# Quantifying the effects of arbuscular mycorrhizal fungi and potato cyst nematodes on root system architecture using X-ray computed tomography

**DOI:** 10.64898/2026.03.09.710487

**Authors:** Eric C. Pereira, Saoirse Tracy

## Abstract

Crop root systems develop in biologically complex soils where beneficial symbionts and pathogenic organisms can jointly influence root architecture and, consequently, belowground function. In this work, we used X-ray computed tomography (CT) to assess how colonisation by the arbuscular mycorrhizal fungus *Rhizophagus irregularis* (AMF) and infection by the potato cyst nematode *Globodera pallida* (PCN) influence root system architecture in soil-grown tomato and potato plants. Root architectural traits, including root volume and root surface area, were quantified non-destructively from intact root systems to evaluate the individual and combined effects of AMF colonisation and PCN infection over time. AMF inoculation increased root volume and surface area, whereas PCN infection caused pronounced reductions in these traits, particularly during early development. AMF-associated increases in root system size were maintained in both PCN-free and PCN-infected plants, indicating largely additive effects of beneficial and pathogenic soil biota on root architectural outcomes. These findings show that soil organisms can independently reshape crop root development in ways likely to influence soil exploration and resource acquisition under biologically complex conditions. More broadly, the study highlights the value of X-ray CT as a non-destructive approach for linking belowground biotic interactions with functionally relevant root traits in sustainable agroecosystems.

## INTRODUCTION

Root system architecture plays a central role in plant performance by regulating water and nutrient acquisition, anchorage, and interactions with the surrounding soil environment (Lynch, 1995; Lynch, 2022; Freschet et al., 2021). Beyond providing structural support, roots define the spatial interface through which plants explore soil resources, and variation in root system organisation strongly influences plant growth, vigour, and resilience under both favourable and stressful conditions. Interactions with the broader root microbiota also play a key role in modulating architecture and stress responses (Balestrini et al., 2024). Despite its importance, quantitative analysis of root architecture remains challenging under soil-grown conditions, where roots are embedded within a heterogeneous matrix and interact dynamically with biotic and abiotic factors. Traditional root phenotyping approaches, including destructive harvesting and washing, disrupt native root–soil relationships and often fail to capture the three-dimensional organisation of intact root systems. Consequently, there is a persistent methodological gap in our ability to quantify root architecture *in situ*.

X-ray computed tomography (CT) has emerged as a powerful non-invasive imaging approach for three-dimensional visualisation of roots growing in soil (Tracy et al., 2010; Mairhofer et al., 2013; Hou et al., 2022). By reconstructing intact soil cores, X-ray CT enables direct observation of the spatial distribution and temporal development of roots without disturbing the soil–root interface (Hou et al., 2022; Ghosh et al., 2023). Importantly, X-ray CT-based methods also enable quantitative extraction of architectural traits (Figure 1), such as root volume and surface area, thereby enabling robust comparisons across treatments and developmental stages (Tracy et al., 2012). While X-ray CT has been widely applied to characterise root growth responses to soil structure and physical constraints, its use for quantifying the effects of interacting soil biota on root system architecture remains comparatively limited (Rogers et al., 2016; Van Harsselaar et al., 2021; Zhang et al., 2022).

**Figure 1.**
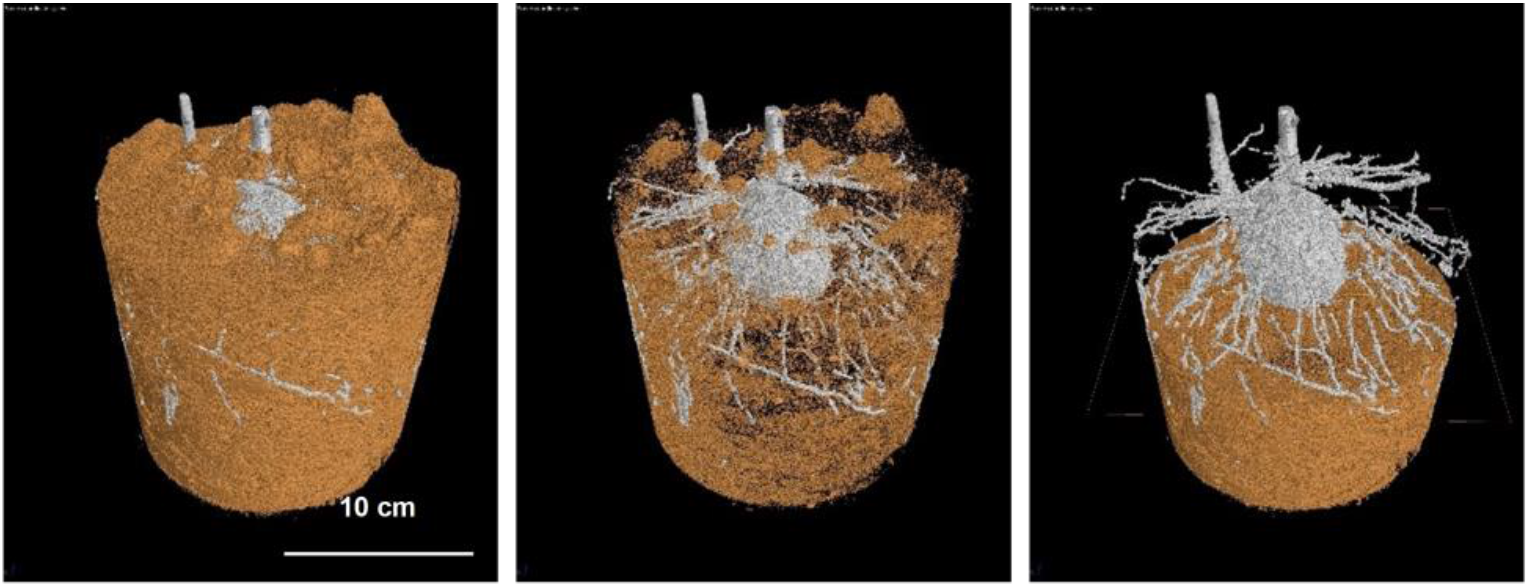
Three-dimensional X-ray computed tomography reconstruction of a potato root system grown in soil, illustrating the spatial distribution of roots (white) within the soil matrix (orange). Scale bar = 10 cm.

Solanum crops, including tomato (*Solanum lycopersicum* L.) and potato (*Solanum tuberosum*), are globally important due to their high nutritional and economic value (Raiola et al., 2014; Jansky et al., 2019). Their production, however, is strongly constrained by soil-borne pests and pathogens. Among these, potato cyst nematodes (PCN) are among the most damaging threats, causing substantial yield losses in potato and other solanaceous crops worldwide (Moens et al., 2018; Price et al., 2021). Species such as *Globodera pallida* are obligate biotrophs that persist in soil as long-lived cysts and impair plant performance by altering root development and nutrient uptake (Turner & Rowe, 2006; De Ruijter & Haverkort, 1999). In contrast, arbuscular mycorrhizal fungi (AMF) form widespread mutualistic associations with Solanum species and contribute to plant nutrition by extending the effective absorptive capacity of the root system, particularly for phosphorus acquisition (Smith et al., 2011; Chen et al., 2018). AMF colonisation has also been shown to influence root system architecture, including changes in root length, branching patterns, surface area, and volume (Chen et al., 2021; Zhang et al., 2021).

Both PCN infection and AMF colonisation are therefore known to modify root development, yet their combined effects on root system architecture within intact soil environments remain poorly quantified. This is an important gap because, in agricultural soils, crop root systems develop within biologically complex environments where mutualists and pests act simultaneously to influence resource capture, stress tolerance, and plant performance. Understanding how beneficial and pathogenic soil organisms jointly shape root system architecture is therefore important not only for root biology but also for predicting crop function under realistic soil conditions and for developing biologically informed management strategies in agroecosystems. Here, we used X-ray computed tomography as a non-destructive phenotyping approach to quantify how colonisation by the arbuscular mycorrhizal fungus *R. irregularis* and infection by the potato cyst nematode *G. pallida* independently and jointly alter root system architecture in soil-grown tomato and potato. By measuring root volume and surface area from intact three-dimensional root systems across developmental stages, we aimed to determine how beneficial and pathogenic soil biota reshape crop root architecture under controlled soil-based conditions and to establish a framework for linking these interactions to crop function in agroecosystems.

## Material and methods

### Plant material and experimental design

Greenhouse experiments were conducted independently for tomato (*Solanum lycopersicum* L.) and potato (*Solanum tuberosum* L.) using a fully factorial design with two factors: arbuscular mycorrhizal fungi (AMF; uninoculated or inoculated) and potato cyst nematode (PCN; non-infected or infected). Each treatment combination was replicated [n = 10] times per species.

Tomato seeds and potato tubers were planted in 60 mL and 2000 mL pots, respectively, containing a sand–soil mixture (1:1, v/v). The substrate was autoclaved at 121 °C for 30 min prior to planting to minimise background microbial activity. AMF inoculation was performed at planting using an inoculum of *Rhizophagus irregularis*. Uninoculated controls received an equivalent volume of sterilised inoculum material to control for substrate effects.

PCN infection was established at planting by incorporating *Globodera pallida* inoculum into the substrate at a density of 50 cysts per pot, determined prior to planting using standard extraction and counting procedures. Control plants received PCN-free substrate. Plants were harvested at 2 and 4 weeks after planting. At each time point, plants were subjected to X-ray computed tomography (CT) scanning prior to destructive sampling for biomass measurements and confirmation of AMF colonisation and PCN infection.

### X-ray CT analysis

To determine the effects of AMF and PCN on the root architecture of tomato plants, all pots were analysed using the GE Nanotom M X-ray CT machine (GE Measurement and Control Solutions). The v|tome|x M was set at a voltage of 65 kV and a current of 300 μA to optimise contrast between background soil and roots. The ‘Fast Scan option’ achieved a voxel resolution of 1.60 μm. 1,078 projection images were taken per scan at 200 m/s per image. Once scanning was complete, the images were reconstructed using Phoenix datos|×2 rec reconstruction software, combining the scans into a single 3D volume representing the entire core.

### Image processing

Image analysis of X-ray CT images was performed using VGStudioMax® (Version 3.2; Volume Graphics GmbH, Heidelberg, Germany) to segment cyst nematodes. Cysts were segmented by setting seed points and using selected threshold values in the Region grower, thereby selecting grey-scale pixels associated with root materials. Once the cysts were segmented from the image, the Erosion and Dilation tool was selected with a 1-pixel radius, and the Region Growing tool was used. Root system architecture parameters, including root length, volume, and surface area, were measured from segmented root systems.

### Detection of AMF and PNC in Inoculated and Infected Plants

Detection of AMF and PCN in plants was conducted by microscopic examination. Root samples were harvested from inoculated and infected plants at harvest to facilitate this analysis. For assessment of fungal structure by microscopy, root specimens were prepared using either the potassium hydroxide (KOH) clearing method or non-clearing methods, followed by staining with Chinese ink or aniline blue, as described by Vierheilig et al. (2005). Conversely, for PCN analysis, cleared root preparations were stained with fuchsin acid, as detailed in the method established in the relevant literature (Byrd et al., 1983).

### Statistical Analysis

The datasets were evaluated for ANOVA assumptions using the Shapiro–Wilk normality test and the Brown–Forsythe test for equal variances. Then, the effects of AMF and PCN on tomato and potato parameters were analysed by two-way ANOVA. Differences between means were evaluated using Tukey’s test. All statistical analyses were performed using SPSS v24.

## RESULTS

### X-ray CT imaging of intact root systems in soil

X-ray computed tomography enabled clear visualisation of intact root systems within the soil matrix for both tomato and potato plants (Figure 2). Three-dimensional reconstructions allowed discrimination between soil, root tissue, and air-filled pores based on grayscale intensity, with roots exhibiting lower X-ray attenuation than the surrounding mineral substrate. These reconstructions provided a basis for consistent segmentation and quantitative analysis of root system architecture without disturbing the soil–root interface.

**Figure 2.**
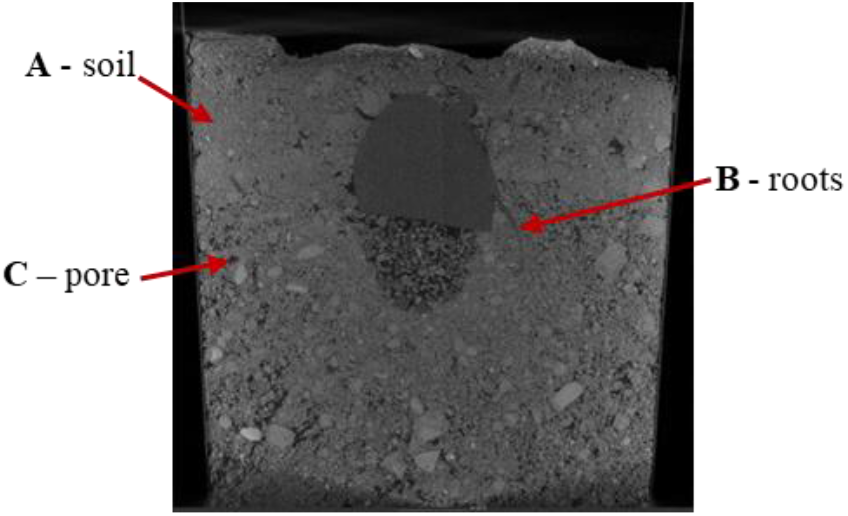
Representative X-ray computed tomography slice of a pot containing a potato plant grown in a soil:sand (1:1, v:v) substrate. Different constituents are visible based on grayscale intensity: (A) soil, (B) root tissue, and (C) air-filled pores. Scale bar = 10 cm.

### Effects of AMF and PCN on root system architecture in tomato

Three-dimensional CT reconstructions revealed clear differences in root system architecture among treatments in tomato plants at both 2 and 4 weeks after planting (Figure 3). Quantitative analysis showed that both AMF inoculation and PCN infection significantly influenced root volume and root surface area, whereas no significant AMF × PCN interaction was detected at either time point (Figure 4; Table 1).

**Table 1.** Results of two-way analysis of variance (ANOVA) testing the effects of AMF inoculation, PCN infection, and their interaction on root volume and root surface area in tomato and potato plants at 2 and 4 weeks after planting.

|  |  |  | AMF inoculation |  | PCN infection |  | AMF x PCN |  |
| --- | --- | --- | --- | --- | --- | --- | --- | --- |
|  |  |  | F | P | F | P | F | P |
| Tomato plants | 2 weeks | Root volume | 3.418 | 0.101 | 7.685 | 0.024 | 1.536 | 0.250 |
|  |  | Root surface area | 2.597 | 0.146 | 8.431 | 0.020 | 1.645 | 0.235 |
|  | 4 weeks | Root volume | 6.024 | 0.040 | 16.83 | 0.003 | 0.413 | 0.538 |
|  |  | Root surface area | 6.120 | 0.038 | 25.16 | 0.001 | 0.309 | 0.593 |
| Potato plants | 2 weeks | Root volume | 36.71 | <0.001 | 8.521 | 0.019 | 0.018 | 0.897 |
|  |  | Root surface area | 12.88 | 0.007 | 11.46 | 0.010 | 0.718 | 0.421 |
|  | 4 weeks | Root volume | 14.93 | 0.005 | 3.461 | 0.099 | 0.011 | 0.918 |
|  |  | Root surface area | 14.95 | 0.005 | 1.678 | 0.231 | 0.954 | 0.357 |

**Figure 3.**
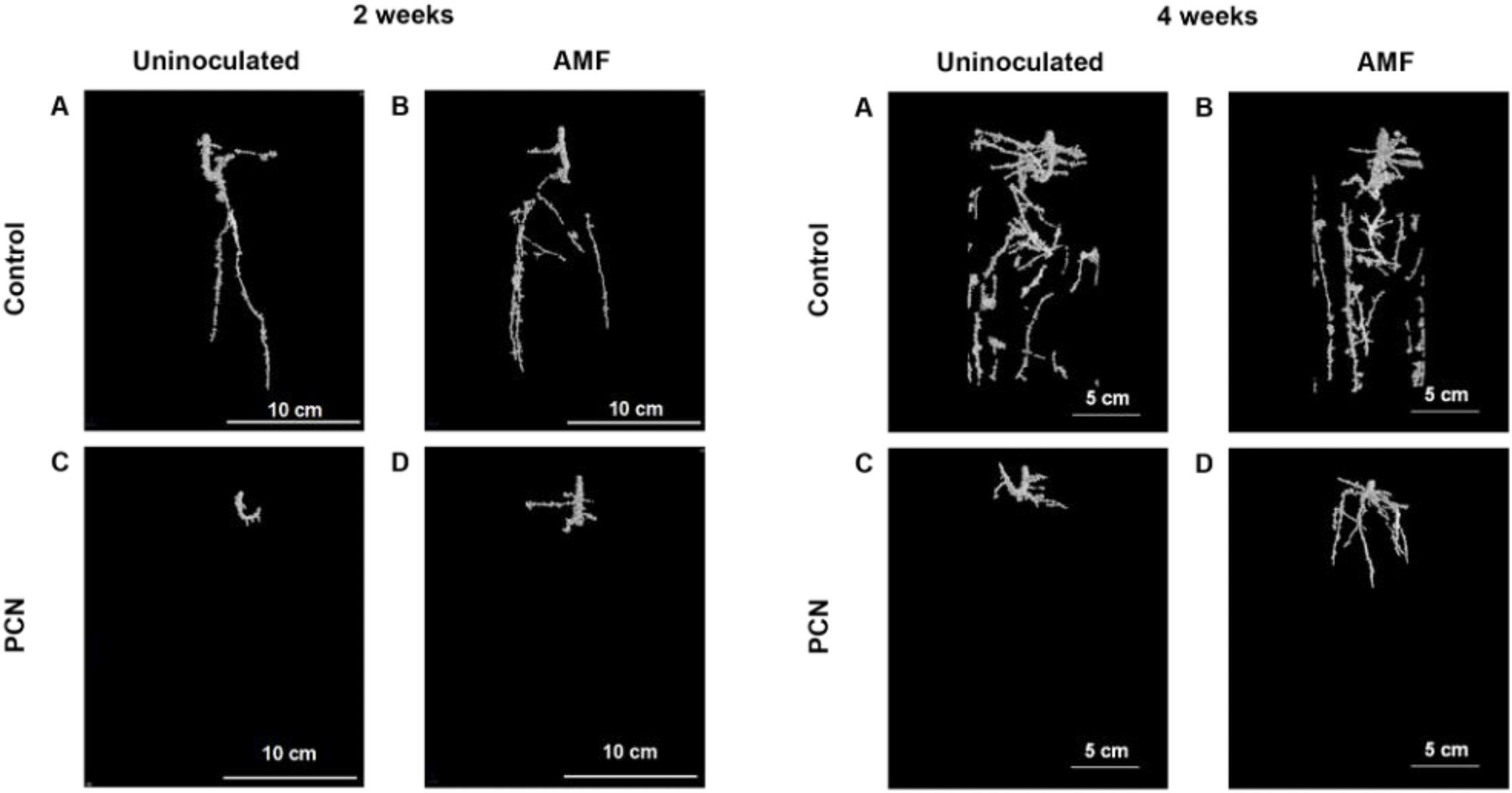
Three-dimensional X-ray computed tomography reconstructions of tomato root systems at 2 and 4 weeks after planting. Roots are shown for uninoculated and AMF-inoculated plants under PCN-free and PCN-infected conditions, illustrating treatment-dependent differences in root system architecture.

**Figure 4.**
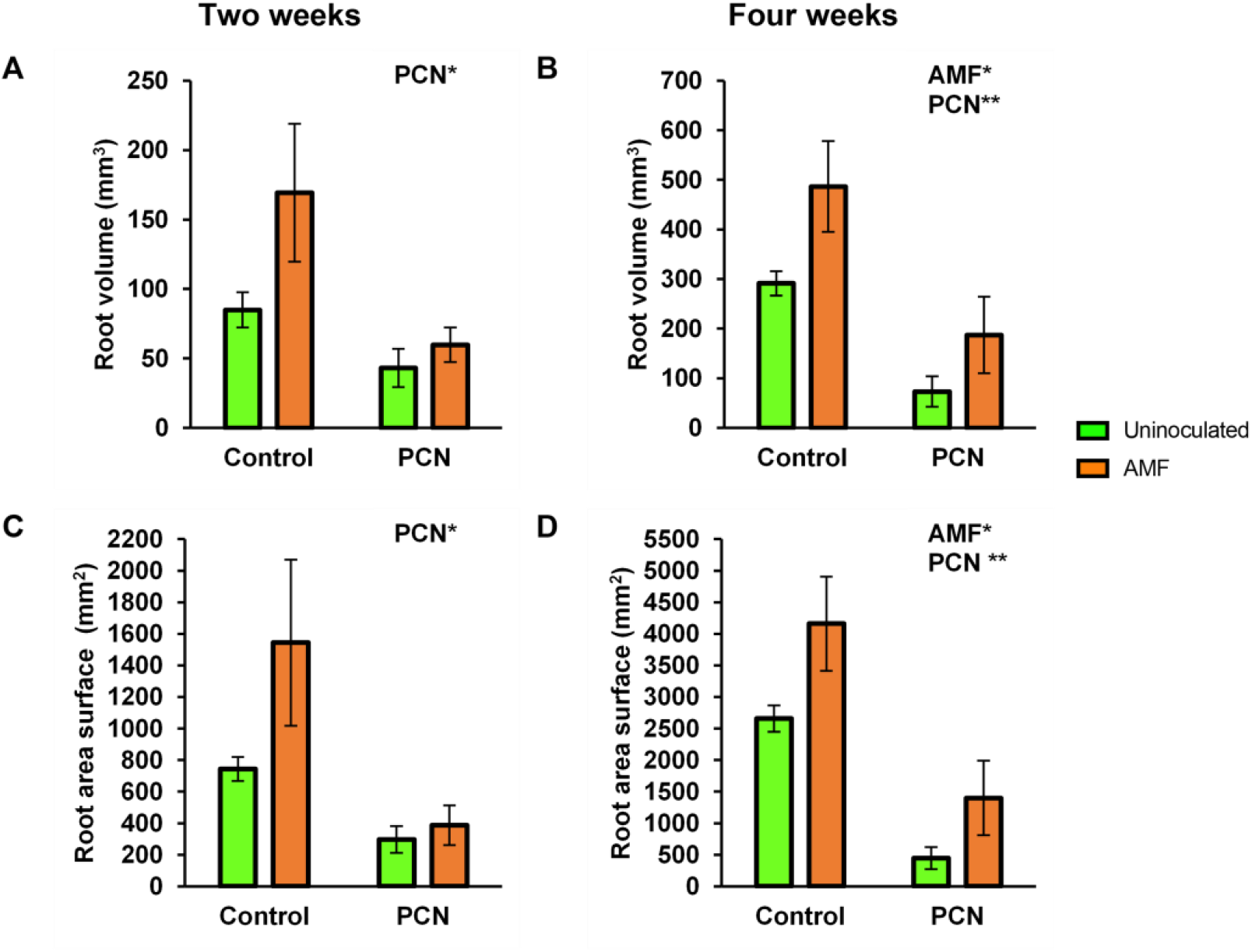
Root system architectural traits of tomato plants quantified by X-ray CT. Root volume (A, B) and root surface area (C, D) are shown for uninoculated (green) and AMF-inoculated (orange) plants under PCN-free and PCN-infected conditions at 2 weeks (A, C) and 4 weeks (B, D) after planting. Values represent means ± SD (n = 5). Asterisks indicate significant differences (*p < 0.05; **p < 0.01).

At two weeks, AMF inoculation resulted in a significant increase in root volume relative to uninoculated plants, both in the absence (+50%) and presence (+28%) of PCN infection (Figure 4A). A similar pattern was observed for root surface area, which increased by 52% in PCN-free plants and by 23% in PCN-infected plants following AMF inoculation (Figure 4C). At four weeks, AMF-associated increases in root volume remained significant, with increases of 40% in PCN-free plants and 60% in PCN-infected plants (Figure 4B). Root surface area also increased significantly at this stage, by 36% in PCN-free plants and 68% in PCN-infected plants (Figure 4D).

PCN infection had a strong negative effect on tomato root architecture at both developmental stages. At two weeks, PCN infection significantly reduced root volume in both uninoculated (−48%) and AMF-inoculated plants (−65%) (Figure 4A). At four weeks, PCN-induced reductions in root volume were even more pronounced, reaching 75% in uninoculated plants and 62% in AMF-inoculated plants (Figure 4B). Similar trends were observed for root surface area, with significant reductions at both time points (Figures 4C and 4D). Despite these reductions, AMF-inoculated plants consistently maintained larger root systems than their uninoculated counterparts under PCN infection.

### Effects of AMF and PCN on root system architecture in potato

X-ray CT imaging also revealed treatment-dependent differences in root system architecture in potato plants at both 2 and 4 weeks (Figure 5). As observed in tomato, both AMF inoculation and PCN infection significantly affected root architectural traits, while their interaction was not significant (Figure 6; Table 1).

**Figure 5.**
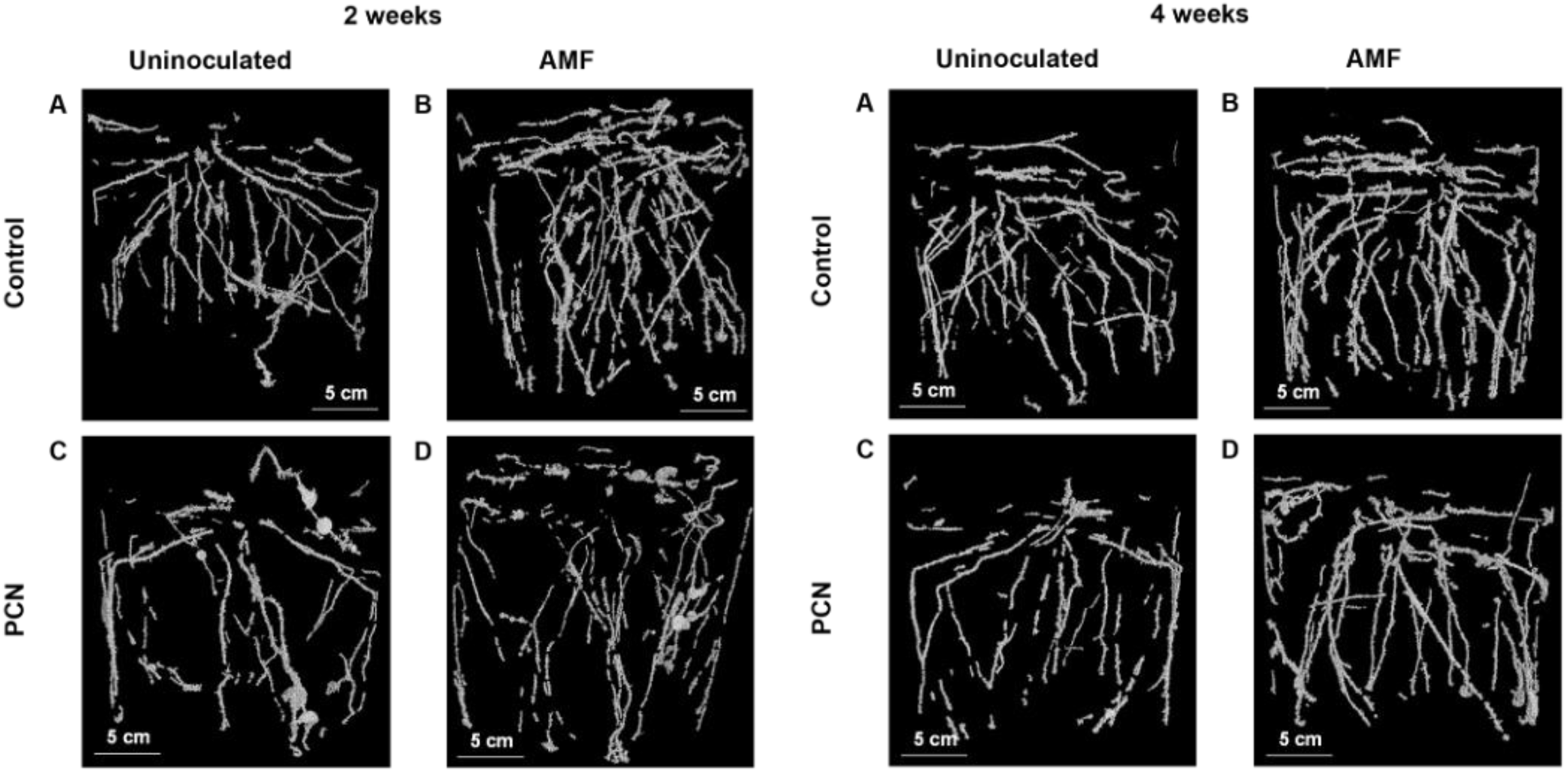
Three-dimensional X-ray computed tomography reconstructions of potato root systems at 2 and 4 weeks after planting. Roots are shown for uninoculated and AMF-inoculated plants under PCN-free and PCN-infected conditions, highlighting treatment-dependent differences in root architecture.

**Figure 6.**
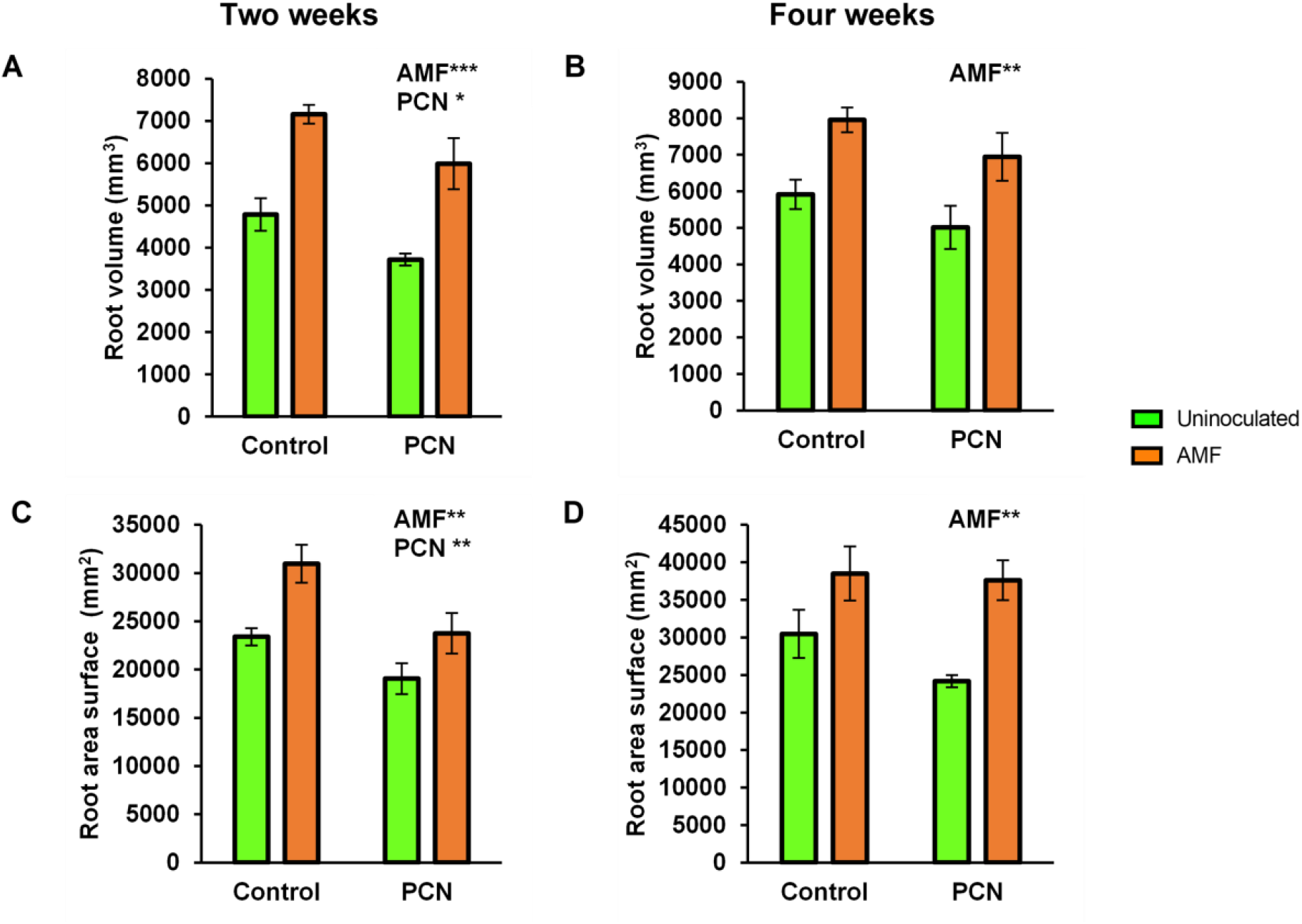
Root system architectural traits of potato plants quantified by X-ray CT. Root volume (A, B) and root surface area (C, D) are shown for uninoculated (green) and AMF-inoculated (orange) plants under PCN-free and PCN-infected conditions at 2 weeks (A, C) and 4 weeks (B, D) after planting. Values represent means ± SD (n = 5). Asterisks indicate significant differences (*p < 0.05; **p < 0.01; ***p < 0.001).

In two-week-old potato plants, AMF inoculation significantly increased root volume by 33% in PCN-free plants and by 38% in PCN-infected plants (Figure 6A). At four weeks, AMF continued to exert a positive effect on root volume, with increases of 26% in PCN-free plants and 28% in PCN-infected plants (Figure 6B). Root surface area responded similarly to AMF inoculation, increasing by 24% (PCN-free) and 20% (PCN-infected) at two weeks (Figure 6C), and by 21% and 36%, respectively, at four weeks (Figure 6D).

PCN infection significantly reduced root volume and surface area in potato plants at the two-week stage. Root volume decreased by 22% in uninoculated plants and by 16% in AMF-inoculated plants under PCN infection (Figure 6A). Root surface area was also significantly reduced, with decreases of 19% in uninoculated plants and 23% in AMF-inoculated plants (Figure 6C). In contrast, PCN effects on root volume and surface area were not statistically significant at four weeks (Figures 6B and 6D).

### Detection of Fungi and PCN in Root Plants

Microscopic examination confirmed the presence of *R. irregularis* structures, including hyphae and spores, in roots of AMF-inoculated tomato and potato plants at both sampling times, irrespective of PCN treatment (Figure 7). Juvenile stages of *G. pallida* were observed in roots of PCN-infected plants, both in the presence and absence of AMF inoculation (Figure 8). These observations confirm the successful establishment of both AMF colonisation and PCN infection in the respective treatments.

**Figure 7.**
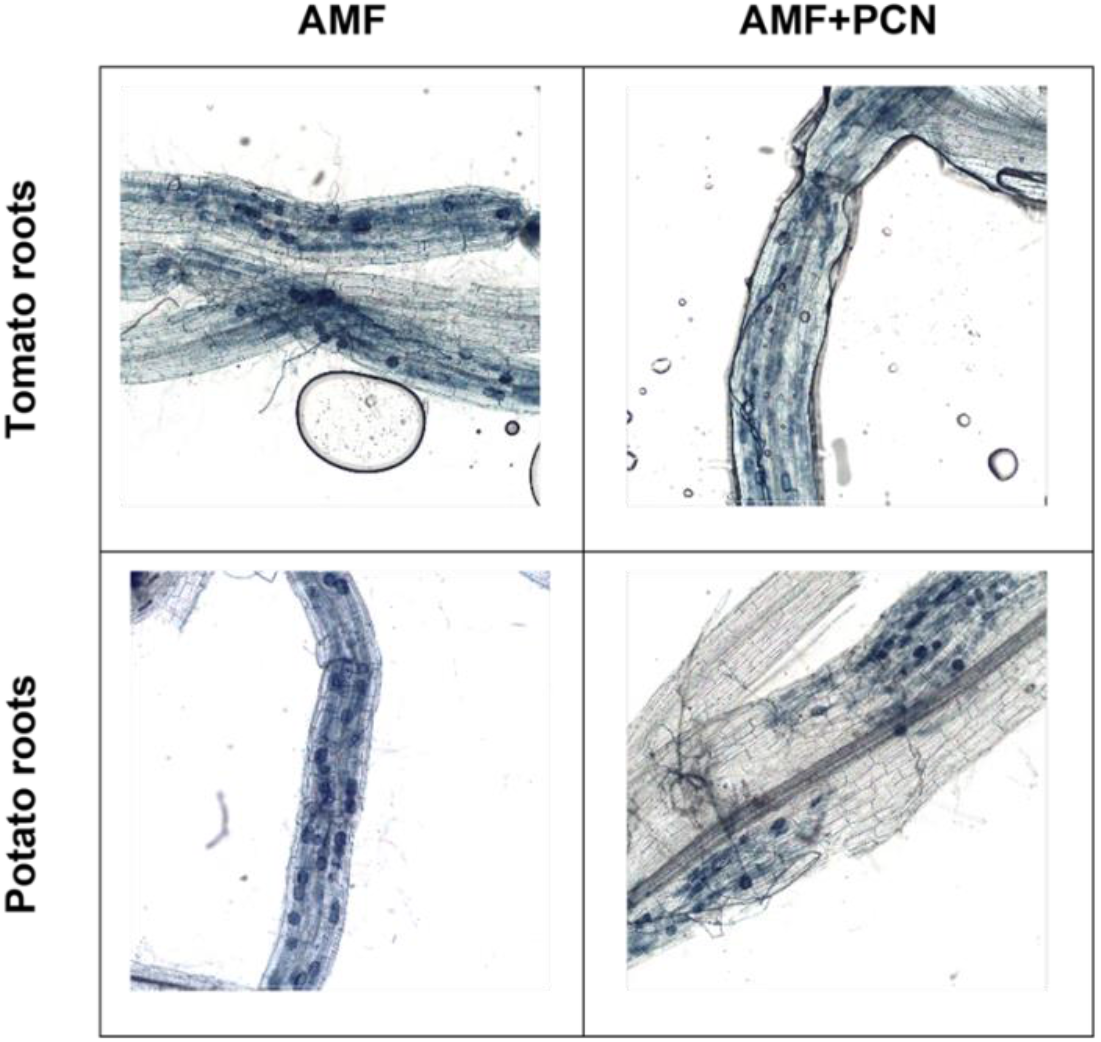
Representative light microscopy images showing *Rhizophagus irregularis* structures in roots of tomato and potato plants inoculated with AMF, including hyphae and spores, under PCN-free and PCN-infected conditions.

**Figure 8.**
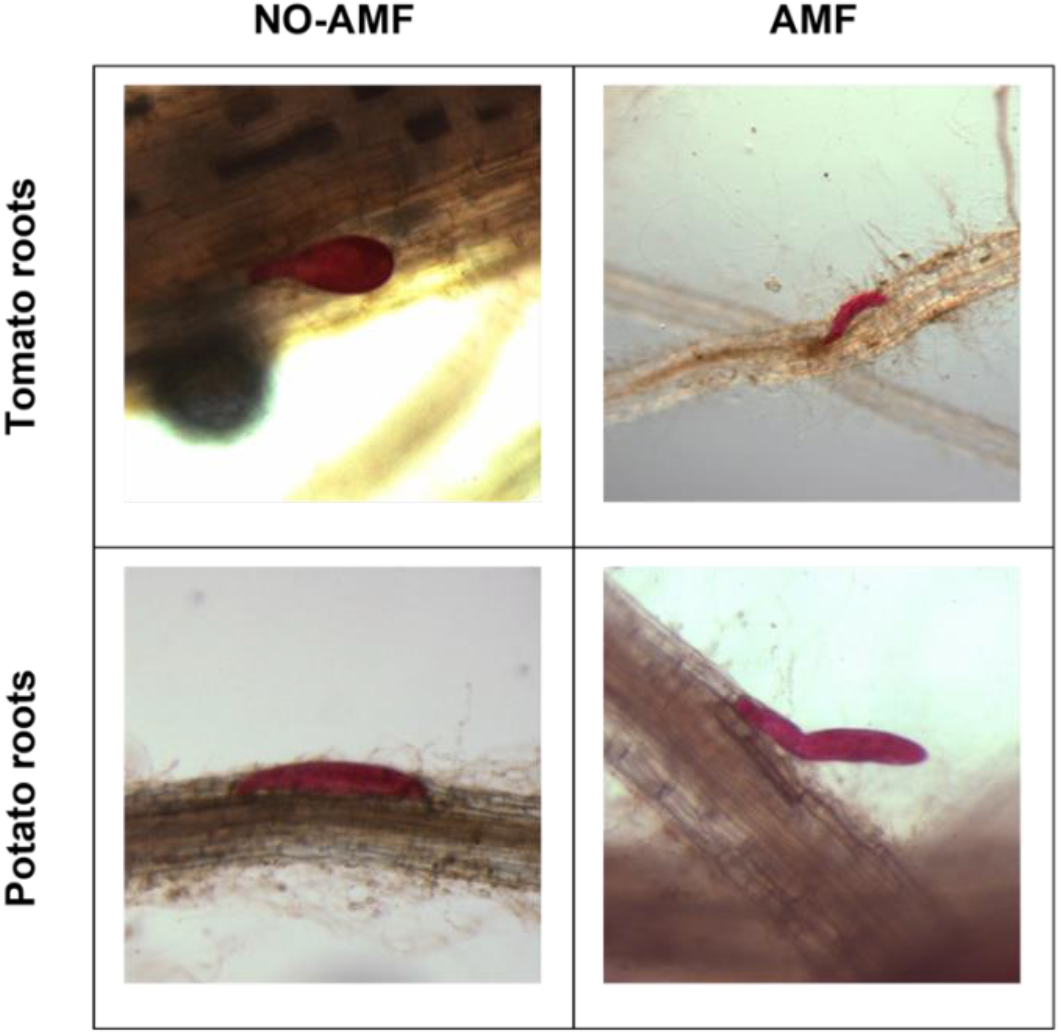
Representative light microscopy images showing juvenile stages of *Globodera pallida* in roots of tomato and potato plants under PCN-infected conditions, in the absence and presence of AMF inoculation.

### Statistical summary

Two-way analysis of variance confirmed significant main effects of AMF inoculation and PCN infection on root volume and root surface area in both species, while AMF × PCN interactions were not significant for any trait or time point (Table 1).

## Discussion

This study shows that beneficial and pathogenic soil organisms can independently and consistently reshape crop root architecture in soil-grown plants. Across tomato and potato, AMF colonisation increased root volume and surface area, whereas PCN infection reduced these traits, particularly during early development. The absence of a significant AMF × PCN interaction indicates that these organisms exerted largely additive effects on root system architecture. These findings are relevant to agroecosystems because root system development underpins soil exploration, resource acquisition, and the capacity of crops to maintain performance under biologically complex soil conditions.

Consistent with previous studies, AMF colonisation by *R. irregularis* was associated with increased root system size, reflected in greater root volume and surface area in both plant species (Begum et al., 2019; Diagne et al., 2020; Ramírez-Flores et al., 2019; Shafiq et al., 2023). Broader microbiome studies show that microbial interactions can drive root phenotypic plasticity (Dini-Andreote et al., 2025). These architectural changes likely reflect AMF-induced modulation of root development, including altered branching patterns and expansion of the absorptive root surface, as reported in earlier work employing destructive phenotyping. In contrast, PCN infection by *G. pallida* exerted a strong negative effect on root system architecture, particularly during early developmental stages. Significant reductions in root volume and surface area were observed in PCN-infected plants, consistent with the well-documented capacity of cyst nematodes to impair root growth and disrupt normal root development (Moens et al., 2018; Palomares-Rius et al., 2017). Together, these results indicate that beneficial and pathogenic belowground organisms can drive contrasting architectural outcomes within the same crop root system.

From an agroecosystem perspective, variation in root system size and spatial development can influence how effectively crops explore soil, intercept water and nutrients, and maintain early vigour under biotic and abiotic stress (Freschet et al., 2021; Lynch, 2022). Increases in root volume and surface area associated with AMF colonisation may therefore indicate enhanced potential for belowground resource capture (Begum et al., 2019), whereas reductions caused by PCN infection likely reflect a constrained capacity to exploit soil resources during early development (De Ruijter & Haverkort, 1999). Although root volume and surface area do not directly measure nutrient uptake or yield, they represent functionally relevant traits through which soil biota may influence crop performance in soil-based production systems (Freschet et al., 2021; Lynch, 2022).

Importantly, although AMF inoculation and PCN infection both significantly influenced root architecture, no significant interaction between these factors was detected for the measured traits. This indicates that AMF-associated increases in root volume and surface area occurred consistently in both PCN-free and PCN-infected plants. Rather than reflecting a specific antagonistic or suppressive interaction between the two organisms, the results point to largely additive effects on root system development. Microscopic observations confirmed the simultaneous presence of AMF structures and PCN juveniles within roots, supporting previous reports that co-colonisation by *R. irregularis* and *G. pallida* does not necessarily result in mutual inhibition (Bell et al., 2022). Within this context, the observed architectural responses represent additive outcomes of symbiotic and pathogenic influences on root system development.

This distinction is important for biologically informed crop management. The present results suggest that AMF-associated changes in root architecture should therefore not be interpreted solely as evidence of direct antagonism against nematodes, but rather as a biologically mediated shift in host root development that persists even when the pest remains present, consistent with broader evidence that mycorrhizal effects on plant defence and performance are mechanistically complex and often host-mediated (Cameron et al., 2013; Vos et al., 2012; Gough et al., 2020). In agroecosystems, such additive outcomes may still be valuable if they help maintain root system function, improve soil exploration, or buffer early developmental damage in infested soils. This is particularly relevant because potato cyst nematodes can impair root growth, nutrient uptake, and crop growth, meaning that partial maintenance of root system development may still have functional benefits even without direct pest suppression (De Ruijter & Haverkort, 1999). More broadly, the findings support the idea that beneficial soil biota may contribute to crop resilience by modulating root traits and host tolerance, even when soil-borne pests remain present (Bell et al., 2022).

Within this broader biological context, X-ray CT provides a valuable means of quantifying how interacting belowground organisms reshape crop root systems in intact soil. Its non-destructive, three-dimensional imaging capacity allows root architectural traits to be measured without disrupting root–soil relationships, thereby preserving the spatial context in which plant–microbe and plant–pathogen interactions occur. The suitability of X-ray CT for plant–nematode systems has previously been supported by the direct detection of potato cyst nematodes in soil-grown plants (Pereira et al., 2025), and the present study extends that framework by demonstrating reproducible treatment effects of AMF colonisation and PCN infection on root volume and surface area in tomato and potato. In this sense, X-ray CT serves as a robust intermediate platform for linking controlled mechanistic studies with more complex agroecosystem questions, enabling more precise investigation of how soil biota influence crop root development under realistic soil-based conditions.

## Conclusion

This study shows that beneficial and pathogenic soil organisms can independently reshape crop root architecture in soil-grown plants. Across tomato and potato, colonisation by *Rhizophagus irregularis* increased root volume and surface area, whereas infection by *Globodera pallida* reduced these traits, particularly during early development. The absence of a significant AMF × PCN interaction indicates that these organisms exert largely additive effects on root system architecture. AMF-associated increases in root system size were maintained even under nematode pressure. Root system size and spatial development influence how effectively crops explore soil, acquire water and nutrients, and maintain function under stress. These findings therefore indicate that soil biota can have important consequences for crop performance in biologically complex soils. X-ray CT was central to this analysis by enabling non-destructive quantification of root architectural traits in intact soil, thereby linking belowground biotic interactions with functionally relevant structural outcomes. More broadly, this study provides a mechanistic foundation for biologically informed strategies to improve crop resilience and manage soil-borne constraints in sustainable agroecosystems.

## Acknowledgements

We gratefully acknowledge funding from the Leverhulme Trust (RPG-2019-162).

## Author contributions

**Conceptualization:** Eric C. Pereira, Saoirse Tracy.

**Data curation:** Eric C. Pereira, Saoirse Tracy.

**Investigation:** Eric C. Pereira.

**Methodology:** Eric C. Pereira, Saoirse Tracy.

**Supervision:** Saoirse Tracy.

**Writing – original draft:** Eric C. Pereira.

**Writing – review & editing:** Eric C. Pereira, Saoirse Tracy.

